# Clickable PEG-norbornene microgels support suspension bioprinting and microvascular assembly

**DOI:** 10.1101/2024.11.15.623424

**Authors:** Irene W. Zhang, Lucia S. Choi, Nicole E. Friend, Atticus J. McCoy, Firaol S. Midekssa, Eben Alsberg, Sasha Cai Lesher-Pérez, Jan P. Stegemann, Brendon M. Baker, Andrew J. Putnam

## Abstract

The development of perfusable and multiscale vascular networks remains one of the largest challenges in tissue engineering. As such, there is a need for the creation of customizable and facile methods to produce robustly vascularized constructs. In this study, secondarily crosslinkable (clickable) poly(ethylene glycol)-norbornene (PEGNB) microbeads were produced and evaluated for their ability to sequentially support suspension bioprinting and microvascular self-assembly towards the aim of engineering hierarchical vasculature. The clickable PEGNB microbead slurry exhibited mechanical behavior suitable for suspension bioprinting of sacrificial bioinks, could be UV crosslinked into a granular construct post-print, and withstood evacuation of the bioink and subsequent perfusion of the patterned void space. Endothelial and stromal cells co-embedded within jammed RGD-modified PEGNB microbead slurries assembled into capillary-scale vasculature after secondary crosslinking of the beads into granular constructs, with endothelial tubules forming within the interstitial space between microbeads and supported by the perivascular association of the stromal cells. Microvascular self-assembly was not impacted by printing sacrificial bioinks into the cell-laden microbead support bath before UV crosslinking. Collectively, these results demonstrate that clickable PEGNB microbeads are a versatile substrate for both suspension printing and microvascular culture and may be the foundation for a promising methodology to engineer hierarchical vasculature.

## 1. Introduction

There is a significant need for functional and perfusable vasculature in the field of tissue engineering, often cited as one of the biggest remaining hurdles to producing functional and clinically-relevant sized artificial tissues and organs.^[1]^ In the absence of functional vasculature, tissue constructs containing metabolically-active cells are limited by the diffusion limits of oxygen to thicknesses of ∼100-200 µm;^[2]^ as a consequence, successful clinical translation of engineered tissues over the past 25 years has largely been restricted to either avascular (e.g., cartilage) or relatively thin tissues (e.g., skin grafts, bladder). To recapitulate tissues with higher complexity, such as solid organs, functional hierarchical vascular trees are needed.^[3]^

While many vascularization strategies focus on the engineering of vessels of a particular length scale (i.e., capillary networks or large blood vessels), the human circulatory system is a complex network of branching vessels with various diameters, wall thicknesses, and wall composition.^[4]^ The combination of larger diameter vessels (allowing for mass transport via advective flow) and capillaries (allowing for diffusive transmural transport) leads to a highly efficient oxygen and nutrient transport system throughout tissue structures. Strategies to engineer hierarchical vasculature have begun to emerge in the literature, and can generally be classified as top-down, bottom-up, or combinations thereof. Top-down, or engineering-driven,^[5]^ strategies typically leverage microfabrication/microfluidics^[6–9]^ or bioprinting^[10–15]^ to template vascular structures and patterns that can be subsequently cultured with endothelial or parenchymal cell populations. Microfabrication strategies are often not amenable to scale-up, while bioprinting techniques typically lack the spatial resolution to engineer capillary-scale networks and have yet to demonstrate a fully functional vascular hierarchy. Bottom-up, or nature-driven,^[5]^ strategies leverage cell-based self-assembly of microvascular networks via culture of endothelial cells in hydrogels either in monoculture or in co-culture with stromal cell populations,^[6, 16–19]^ wherein the latter, stromal cells support the development, maturation, and stabilization of the vessel networks and adopt a perivascular phenotype.^[20–22]^ Promising recent methods to engineer multi-scale vasculature have combined both top-down and bottom-up strategies. For example, tissue-engineered vascular grafts fabricated to replicate large-diameter vessels have been embedded in cell-laden hydrogels to produce multi-scale vascular structures.^[23–24]^

Our approach to partially address this persistent vascularization challenge likewise aims to use a combination of top-down and bottom-up strategies, combining suspension bioprinting of mesoscale vascular patterns with cell-initiated microvascular self-assembly to produce vessel-like structures of different sizes. Suspension baths are often temporary, intended to be dissolved, degraded, or otherwise liquified to release the printed structure after it has solidified.^[15, 25]^ By contrast, suspension baths can also be secondarily crosslinked and retained,^[26–27]^ allowing for customization of the culture environment prior to cell seeding by tuning the composition of the granular bath or by patterning void spaces. Previous studies have demonstrated the ability to template vascular networks via suspension bioprinting of sacrificial inks into supportive granular baths.^[10, 15]^ This latter approach, which is less common in the biofabrication literature,^[28]^ leverages the potential of granular hydrogels, an emerging class of innovative biomaterials formed by physically jamming colloidal hydrogel-based microgels (or microbeads) into microporous monolithic constructs resembling packed beds.^[29]^ These granular materials have useful mechanical properties prior to crosslinking, including shear-thinning and self-healing properties amenable to injectable delivery and bioprinting applications.^[26, 30]^ The microgel building blocks can be linked via non-degradable covalent interactions,^[31]^ reversible non-covalent interactions (e.g., guest-host)^[32–33]^ or proteolytically-susceptible peptide crosslinks to create microporous annealed particle (MAP) scaffolds, which have shown favorable pro-regenerative healing characteristics when applied to chronic skin wounds.^[34–35]^ By tuning the size, shape (e.g., spheres, rods, etc.) and physical properties of the microgel building blocks, the interstitial porosity of granular hydrogels can be designed and controlled to support cell spreading,^[31, 36]^ enable cellular migration, and facilitate mass transport.^[37]^ The facile properties of granular materials have led to their increasing popularity in tissue engineering studies. However, relatively few studies have aimed to vascularize granular materials.^[38–40]^

In this study, we developed clickable PEGNB microbeads and evaluated their ability to support suspension printing of sacrificial bioinks, form granular constructs upon secondary crosslinking after printing, and enable microvascular self-assembly within the interstitial voids of printed constructs. The jammed slurries of the clickable microbeads demonstrated mechanical properties favorable for suspension printing. Sacrificial bioinks were printed in the bead bath, followed by secondary crosslinking to form a free-standing granular construct capable of withstanding manipulation, submersion, evacuation of the sacrificial ink, and perfusion of the patterned void space. Endothelial cells and supportive stromal cells co-embedded within the bead slurries prior to printing self-assembled into mature microvascular networks after both printing and secondary cross-linking of the microbeads into granular constructs, vascularizing a >500 mm^3^ volume construct within 7 days of culture. Together, the data presented in this work demonstrate the use of clickable PEG-based granular materials to sequentially support both suspension bioprinting and microvascular self-assembly in the same construct. This innovative approach enables the production of large, vascularized constructs, and provides a path towards engineering hierarchically vascularized tissues.

## 2. Results

### 2.1. Formation of clickable PEGNB hydrogel microbeads and granular hydrogel constructs

PEGNB-based hydrogel microbeads were formed via thiol-ene UV polymerization with non-degradable crosslinker (PEGDT) and adhesive peptide (RGD) for integrin binding (**Figure 1A**). A crosslinking ratio (% norbornene arms crosslinked, or thiol:ene) of 37.5% (0.375:1 thiol:ene after accounting for RGD concentration) was selected to allow for sufficient remaining functional groups post-microgel polymerization for secondary crosslinking (clicking) with other beads. Beads were formed via microfluidic droplet generation in a flow-focusing device, isolated from oil, and equilibrated in PBS overnight (**Figure 1B**). After swelling, beads were jammed via on-strainer vacuum filtration to form a bead slurry, and subsequently used for rheology, bioprinting, and cell culture (**Figure 1B, C)**. Prior to swelling, the PEGNB beads were analyzed to have an average diameter of 202.09 ± 40.39 µm (**Figure 1D**); after swelling, the beads had an average diameter of 370.37 ± 73.29 µm (**Figure 1D**). The microbeads were highly monodisperse (polydispersity index, PDI < 0.1)^[41]^) across multiple batches, with a PDI of 0.04.

**Figure 1.**
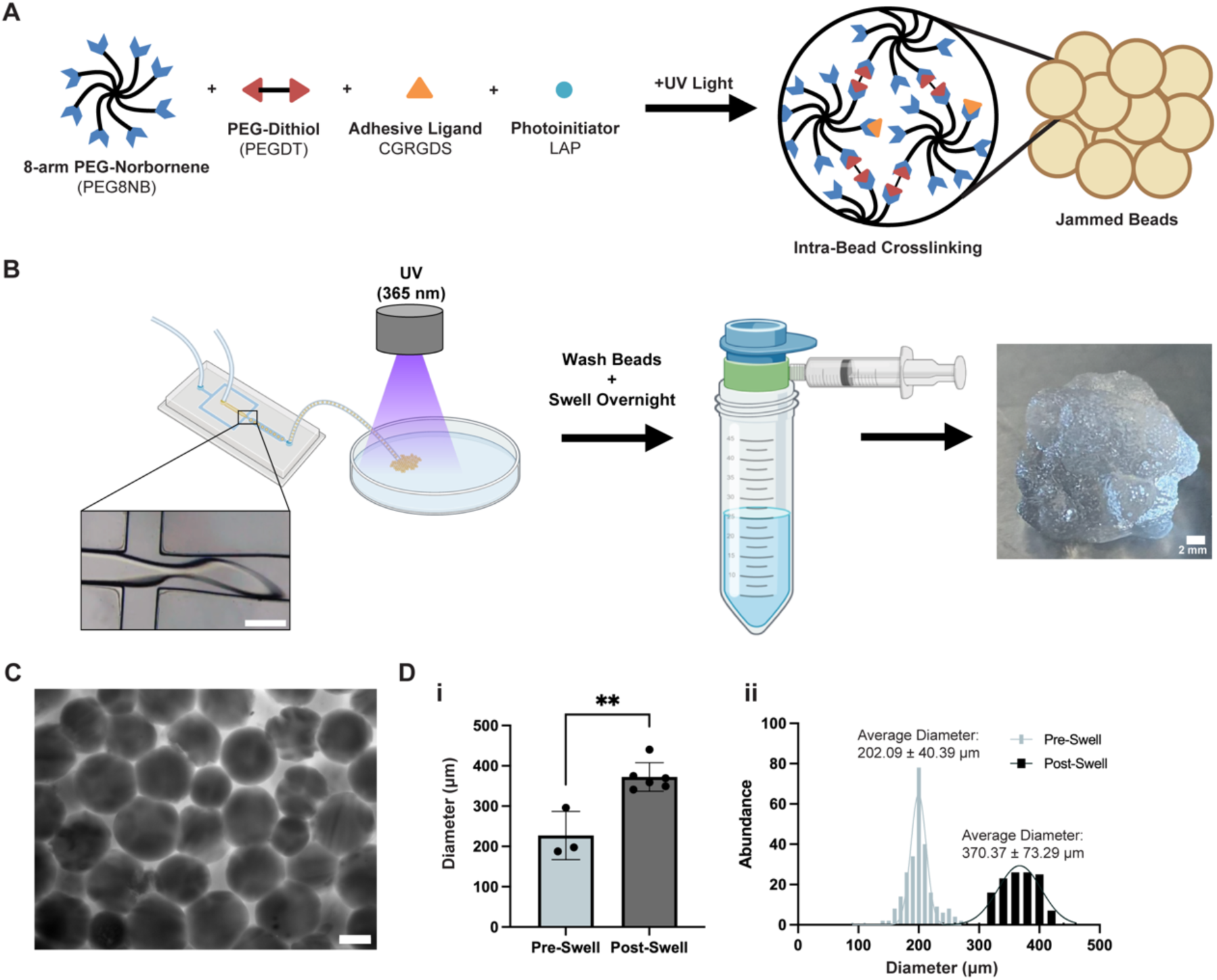
PEGNB microbeads were reliably formed via microfluidic droplet generation. A) Schematic of components and thiol-ene photopolymerization of PEGNB clickable beads. B) Schematic of workflow to produce PEGNB beads and prepare them for experimental use (microfluidic device scale bar = 200 μm). C) Representative 4X magnification image of 10% PEG8NB 37.5% crosslinked 2 mM RGD beads incubated with 5 mg/mL TRITC-dextran (scale bar = 200 μm). D) i) The average diameter of beads per batch before (n = 3) and after (n = 6) overnight swelling in 1X PBS as analyzed via Fiji (p = 0.0023) and ii) distribution of individual bead diameter over multiple batches (beads analyzed per batch ≥ 43; pre-swell, n = 3 batches; post-swell, n = 6 batches).

After the beads were jammed, they could be packed into a PDMS mold and clicked together to form granular hydrogel constructs via the addition of LAP and PEGDT and subsequent UV photopolymerization (**Figure 2A, B)**. Covalent photocrosslinking of the beads was confirmed through *in situ* dynamic rheology. The shear storage modulus (G’, Pa) reached its peak and plateaued after 26 seconds of UV exposure at 13.2 mW/cm^2^ (**Figure 2C**). This increase in G’ was maintained over time, indicating that crosslinking of the beads formed a stable, free-standing granular construct. To visualize the interconnected pore space between beads, PEGNB beads made with thiolated rhodamine B were formed into clicked granular constructs and incubated with high molecular weight FITC-dextran (2 MDa) and imaged via confocal microscopy (**Figure 2D, Movie S1)**. Average percent porosity per imaged slice was 23.4 ± 5.5%.

**Figure 2.**
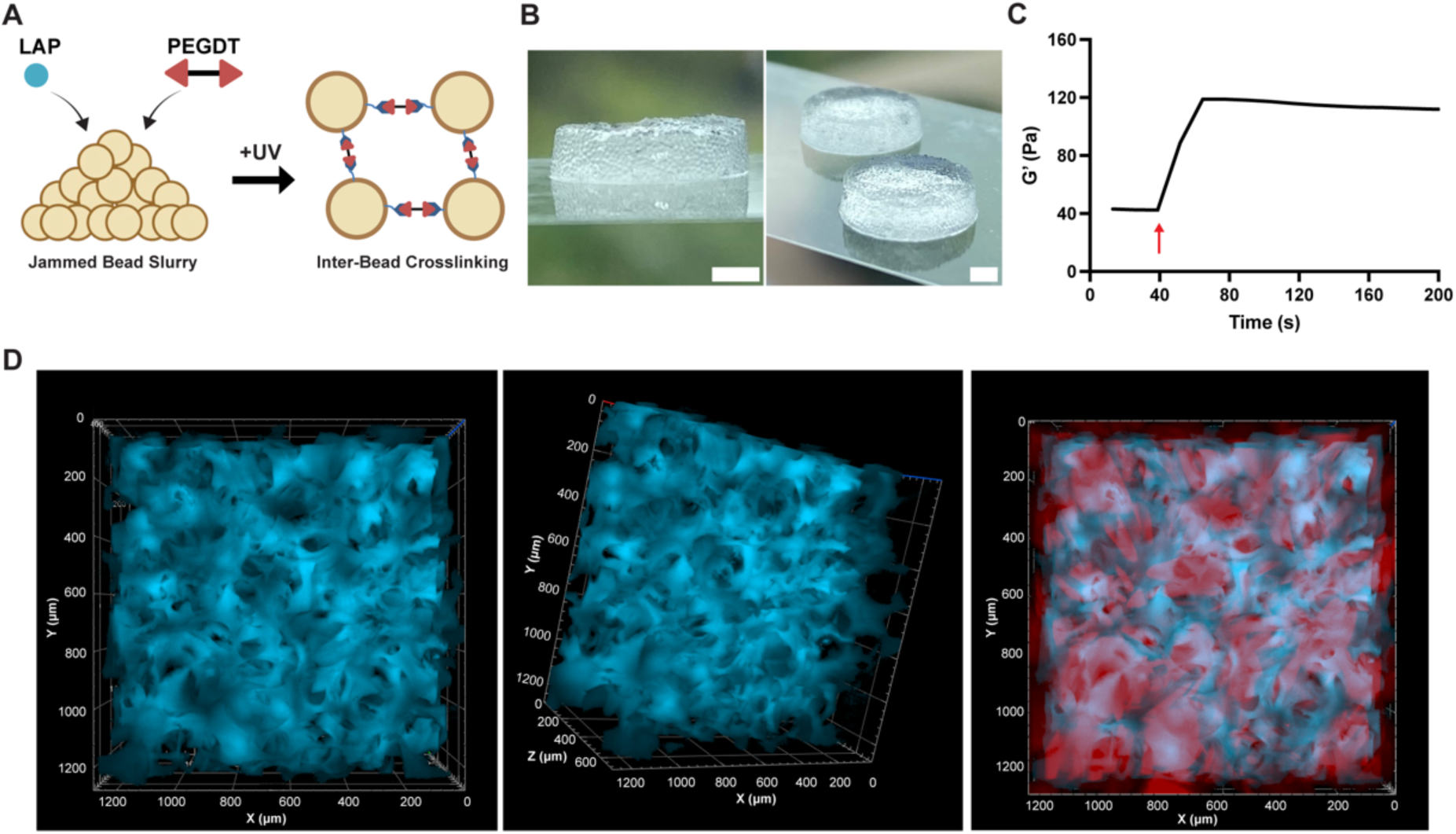
PEGNB microbeads were photopolymerized into covalently crosslinked (clicked) granular constructs. A) Schematic of PEGNB jammed slurry photopolymerization into clicked granular construct. B) Images of clicked granular constructs (scale bar = 2 mm). C) Representative time-sweep trace of the shear storage modulus (G’) of bead slurry with additional LAP and PEGDT pre- and post-UV exposure (arrow indicating UV turned on, exposed from t = 40 s to t = 200 s) at an intensity of 13.2 mW/cm^2^. D) A representative 3D reconstruction of a clicked granular construct and its void space (blue = void, red = beads).

### 2.2. Clickable PEGNB beads are a suitable medium for suspension bath bioprinting

PEGNB beads were assessed mechanically to determine their suitability for suspension bioprinting (**Figure 3A**). Decreasing viscosity of the PEG microbead slurry with increasing shear rate indicated shear-thinning qualities (**Figure 3B**). An oscillatory strain sweep demonstrated G’ (shear storage modulus) to be greater than G” (shear loss modulus) with increasing strain magnitude for the slurry, until reaching the critical strain (∼66% strain) where the two curves intercepted each other (**Figure 3C**), indicating strain yielding behavior. Lastly, the jammed bead slurry was subjected to alternating low (1% strain) and high (150%) strain. At low strain, the G’ was greater than G”, with the reverse behavior seen at high strain indicating a reversible solid-like and fluid-like behavior integral to suspension bioprinting (**Figure 3D**). This behavior and the magnitude of G’ and G” did not change over successive alternating cycles of low and high strain.

**Figure 3.**
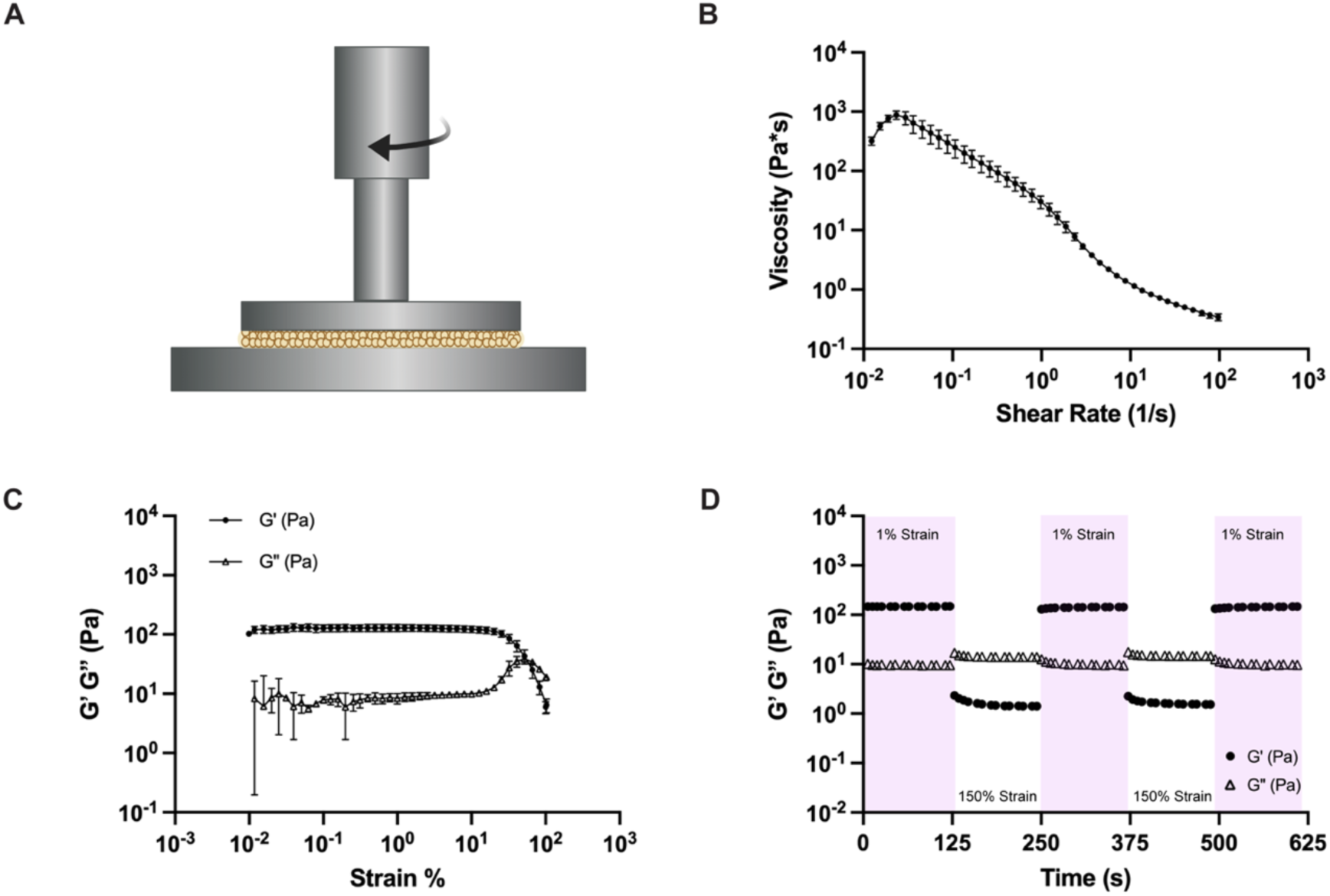
PEGNB bead slurry exhibits mechanical properties suitable for suspension bath printing. A) Schematic of rheometer set-up with bead slurry. B) Rheological characterization demonstrating decreasing viscosity with increasing shear rate (0.01 to 100 1/s) (n = 3), C) strain-yielding behavior with increasing strain (0.01 to 100%) (n = 3), and D) reversible solid-like and fluid-like behavior with low (1%, 1 Hz, purple shaded) and high (150%, 1 Hz, unshaded) strain cycles. A representative trace is shown for time sweep data.

The suitability of the bead slurry for suspension bioprinting was confirmed by printing various structures using an Allevi-3 bioprinter. The hydrogel beads were jammed and packed into a 16 mm diameter x 14 mm height (**Figure 4A**) or 20 mm diameter x 15 mm height (**Figure 4B**) PDMS mold. Clickable PEGNB beads were able to support printing of both gelatin and Pluronic F-127 inks as demonstrated by the cylindrical structures suspended in the center of the bead bath (**Figure 4A**). The bead slurry also supported complex structures that would otherwise collapse if printed in air (**Figure 4B, Movie S2)**. When LAP and PEGDT were added to the bead slurry prior to printing and the bath was exposed to UV post-printing, inter-bead crosslinking occurred, yielding a free-standing granular construct containing the embedded print (**Figure 4B**). Clicked granular constructs were mechanically stable and withstood manipulation despite embedded prints disrupting some inter-bead crosslinking. Granular constructs with embedded gelatin structures withstood submersion in warm PBS and subsequent evacuation of the gelatin ink (**Figure 4C**) without dissociation. This left a patterned void within the granular construct, which could then be perfused (**Figure 4D-F, Movie S3, S4)**. Perfusion of food coloring-dyed glycerol into the patterned void in the granular construct was accompanied by ejection of PBS from the construct due to liquid displacement, which did not affect the overall integrity of the granular hydrogel. Printed structures and subsequent patterned evacuated void spaces measured at diameters ranging between 600 μm to 1 mm, which is within the range of mesoscale vascular formations (50 – 1000 μm).^[3]^

**Figure 4.**
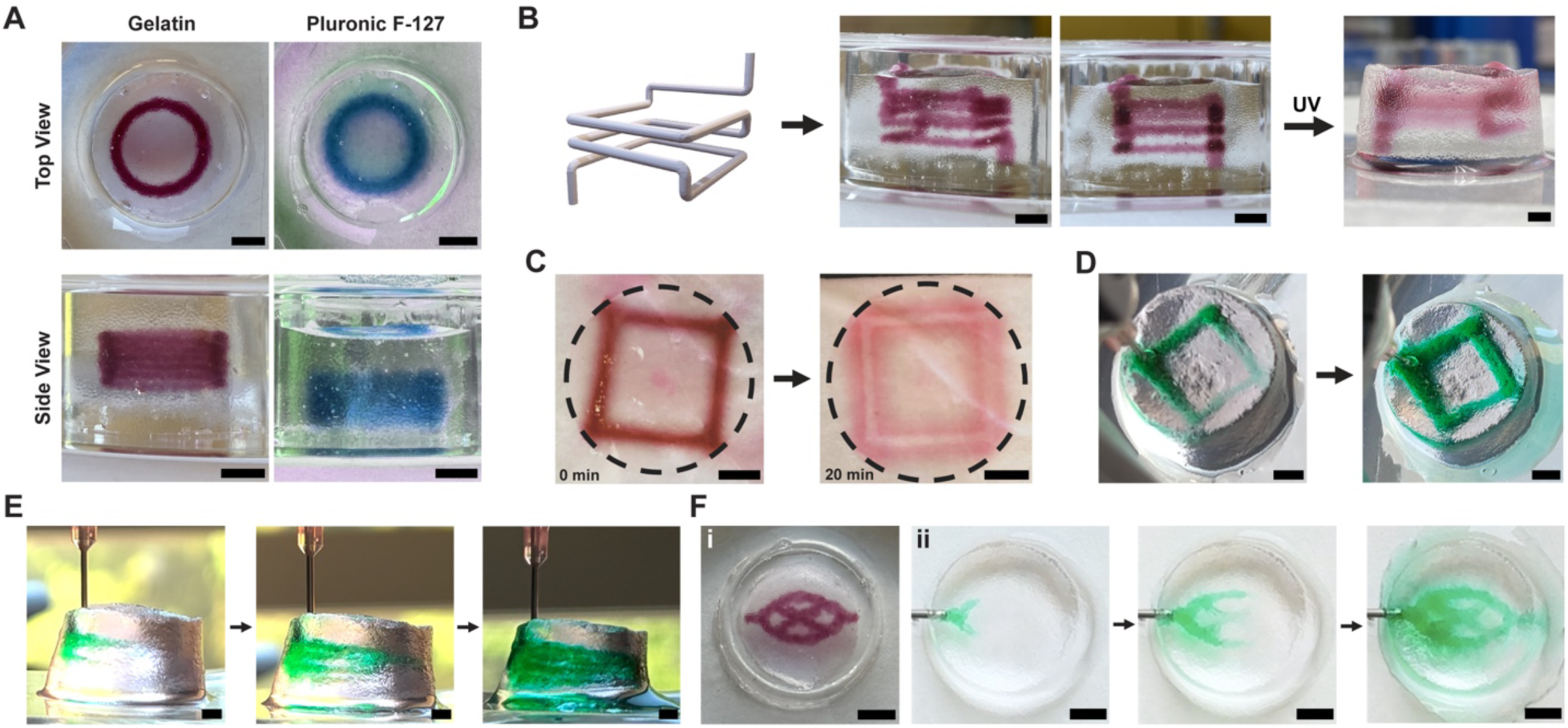
PEGNB bead slurry supports suspension bioprinting and crosslinking into a manipulatable clicked granular construct. A) Representative images of gelatin (dark red) and Pluronic F-127 (blue) bioinks printed to form a cylinder structure and suspended in jammed bead slurry baths in PDMS molds. B) The printing process of a complex, suspended structure into a bead bath, starting with designing a CAD model, to printing the structure with gelatin into the bead bath, to UV crosslinking of the bead bath with the embedded print. C) A clicked PEGNB granular construct with an embedded gelatin print in warmed PBS, allowing for evacuation of the bioink after 20 minutes. D) Void space left from printing CAD structure (panel B) perfused with glycerol (green) from top view and E) side view. F) Top view of capillary-bed like structure i) printed with gelatin (dark red) and ii) perfused with glycerol (green) post-UV and ink evacuation. Scale bar = 4 mm.

### 2.3. Clicked PEGNB granular constructs support microvascular self-assembly

Human umbilical vein endothelial cells (ECs) and normal human lung fibroblasts (LFs) were co-cultured within the clicked PEGNB-based granular constructs to assess their ability to support microvascular self-assembly within the inter-bead void space. LAP and PEGDT were mixed with the jammed bead slurry before adding a suspension of ECs and LFs (referred to as EC-LF when co-cultured) to the mixture. Cells were briefly mixed with the beads before UV photopolymerization to click the beads together (**Figure 5A**). The granular, cell-laden constructs had a volume of 502.65 mm^3^, measured at 16 mm in diameter and 2.5 mm in thickness. To determine optimal cell spreading conditions when cultured in the granular hydrogels, concentrations of secondary LAP and PEGDT were varied alongside cell density. Higher concentrations of LAP and PEGDT were associated with rounded cells and minimal spreading of both cell populations **(Figure S1)**. As LAP and PEGDT concentration decreased and cell density increased (5×10^5^ cells/mL to 1×10^6^ cells/mL), cell morphology of both ECs and LFs changed to exhibit elongation and spreading around beads **(Figure S1)**. The volume of medium also impacted cellular assembly, with ECs exhibiting a more elongated and branching morphology and LFs exhibiting improved spreading when the medium was increased from 1 to 2 mL **(Figure S2)**.

**Figure 5.**
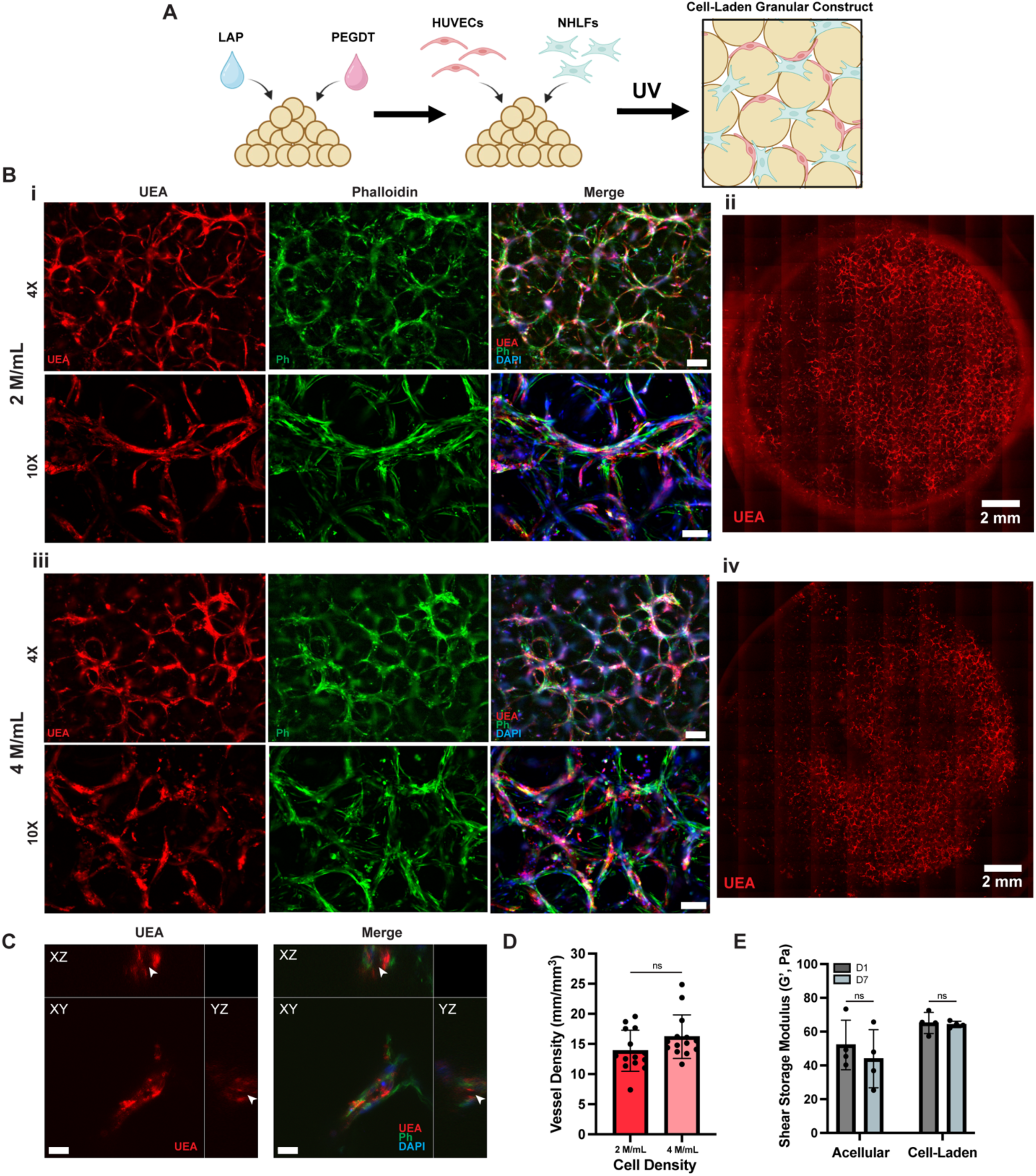
PEGNB granular constructs seeded with ECs and LFs are supportive of microvascular-like self-assembly. A) Schematic of producing cell-laden granular constructs. B) Representative images of vascular development in 2×10^6^ cells/mL (2 M/mL, i, ii) and 4×10^6^ cells/mL (4 M/mL, iii, iv) EC-LF-laden granular constructs after 7 days of culture at 4X (top, scale bar = 200 µm) and 10X (bottom, scale bar = 100 µm). Representative 4X scan-slide images (ii, iv) of the entire EC-LF-laden granular constructs are also shown. C) Orthogonal view of vascular structures in 2×10^6^ cells/mL granular constructs, with arrows denoting lumen formation (scale bar = 20 µm). D) Quantified vessel density of 2×10^6^ cells/mL (n = 14) and 4×10^6^ cells/mL (n = 14) EC-LF-laden granular constructs at day 7 of culture. E) Shear storage modulus on day 1 (n = 4) and day 7 (n = 4) of culture for acellular and cell-laden granular constructs.

Two different cell densities (2×10^6^ cells/mL and 4×10^6^ cells/mL) were examined to evaluate microvascular self-assembly in the granular constructs. After 7 days of culture, constructs exhibited microvascular self-assembly in the inter-bead void space in scaffolds for both cell densities. Cells spread along the surface of the nondegradable RGD-modified microbeads, with ECs (UEA+ cells) forming tubular structures throughout the granular construct (**Figure 5B**). For both cell density conditions, extended EC (UEA+ cells) networks were observed, while LFs (F-actin+, UEA-cells) supported EC morphogenesis in a pericyte-like manner, as demonstrated by their co-localization to UEA-positive vascular structures (**Figure 5B-i, 5B-iii, Figure S3A)**. Lumen formation was confirmed in these vascular structures via orthogonal confocal views (**Figure 5C, Movie S5, S6)**. No significant difference in vascular density at day 7 was observed between 2×10^6^ cells/mL and 4×10^6^ cells/mL seeding conditions, yielding average densities of 13.88 ± 3.41 and 16.21 ± 3.62 mm/mm^3^, respectively (**Figure 5D**). When assessing mechanical stability of the constructs through culture, both acellular and cell-laden granular constructs exhibited no significant changes in shear storage modulus between days one and seven of culture (**Figure 5E**).

To validate the pericyte-like behavior of the LFs further, constructs were stained with antibodies for α-SMA, PDGFR-β, and NG2, known pericyte markers **(Figure S3B)**. The expression patterns of these markers confirmed the close perivascular association of the LFs with the vessel-like structures, suggesting these cells (or a subset of them) are capable of becoming bona fide pericytes **(Figure S3, Figure 5B-i, Figure 5B-iii)**. Positive staining for both NG2 and PDGFR-β showed a sheath-like morphology around microvascular networks **(Figure S3C)**. Constructs were also stained for laminin-β 1 and collagen IV, key components of the basement membrane. Both were found to be closely localized to the UEA-positive vascular formations **(Figure S4)**.

### 2.4. Cell-laden granular constructs maintain the ability to support microvascular self-assembly post-printing and ink evacuation

Next, we aimed to assess whether cell-laden bead baths could withstand mechanical shear forces caused by the disturbance of printing and still support microvascular self-assembly. Cell-laden granular bead slurries were prepared first and then used to support printing of gelatin or Pluronic F-127 cylinders, after which they were UV crosslinked to form constructs that could be cultured (**Figure 6A, B)**. Bioinks evacuated passively over the course of culture time due to incubation at 37 °C and/or media changes. Viability was assessed via live/dead assay on day 1 and 7 of culture **(Figure S5A-B)**. The average percent viability was then normalized to the average of the control (no print) condition (**Figure 6C**) and exhibited no significant differences amongst the three conditions on day one of culture. Printing did not influence cell viability in the extrusion path of the printed structure as well, with the gelatin printed construct even displaying more viable cells at the border of the print **(Figure S5A)**. Viability was also assessed on cell-laden bulk PEG gels made with protease-sensitive peptide crosslinkers^[18]^ on day 1 of culture **(Figure S5C)**. When normalized to the average viability in these bulk PEG gel controls, granular constructs exhibited lower viability (bulk average = 100 ± 0.04%, no print granular = 86.28 ± 0.05%, gelatin print granular = 88.59 ± 0.05%, Pluronic F-127 print granular = 88.55 ± 0.05%), though this may be due to many factors, such as size of construct and LAP concentration used **(Figure S5D)**.

**Figure 6.**
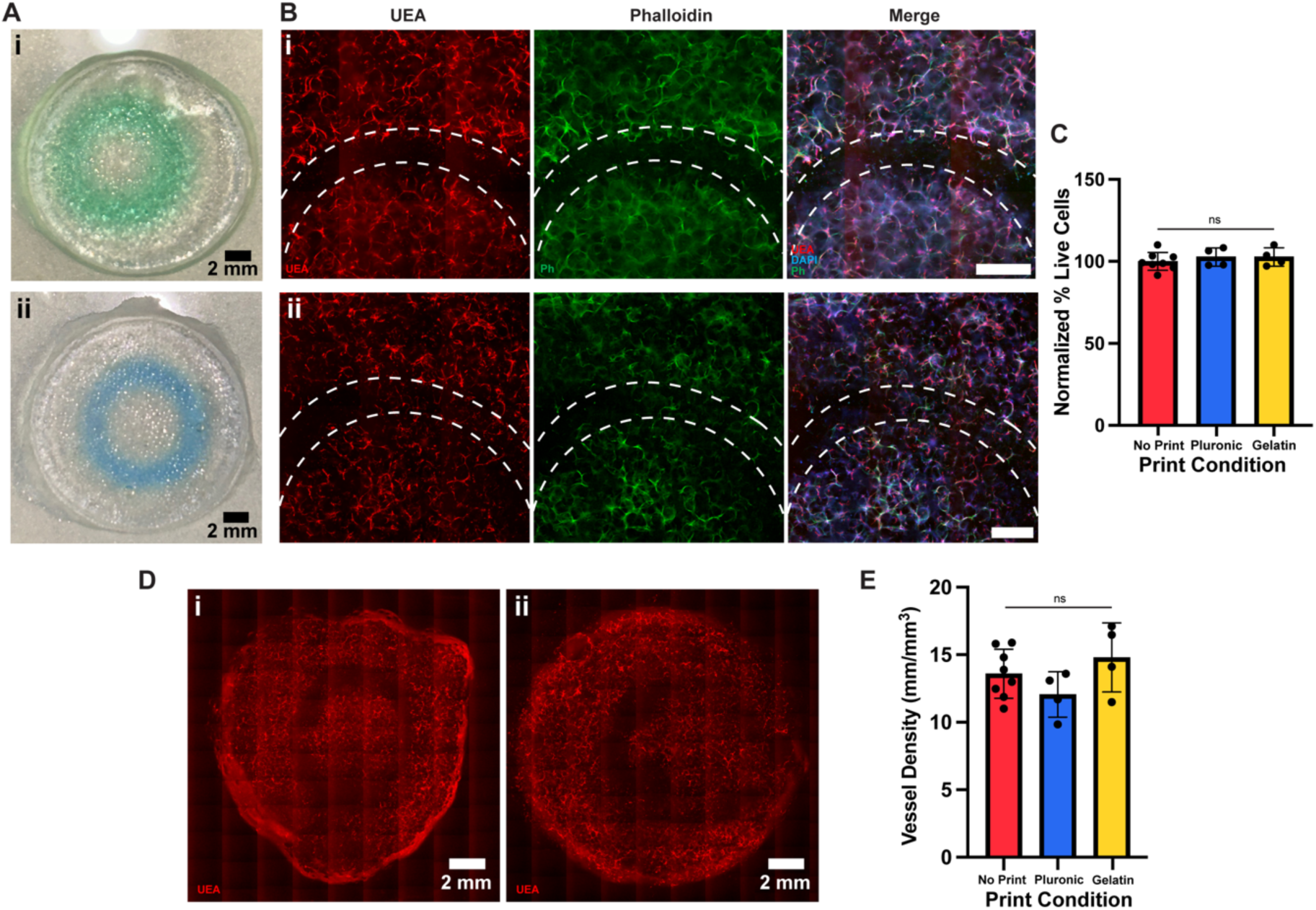
Bioprinting of sacrificial ink into cell-laden PEGNB microbead bath followed by inter-bead crosslinking does not adversely affect microvascular self-assembly. A) Images of cell-laden granular construct with i) gelatin (green) and ii) Pluronic F-127 (blue) cylinder printed and embedded within. B) Representative 4X scan-slide images of i) gelatin-printed and ii) Pluronic F-127-printed cell-laden granular constructs after 7 days of culture (UEA = red, phalloidin = green, DAPI = blue). Dotted line region indicates path of printing. Scale bar = 1 mm. C) Percent live cells in cell-laden granular constructs on day 1 of culture that were printed into with gelatin (n = 4) or Pluronic F-127 (n = 4), normalized to average viability of no print constructs (n = 8) on day 1. D) Representative 4X scan-slide images of vascular development after 7 days of culture in granular constructs which were printed into with i) gelatin or ii) Pluronic F-127 (UEA = red). E) Quantified vessel density in granular constructs with gelatin (n = 4), Pluronic F-127 printing (n = 4), or without printing (n = 8).

Constructs cultured for seven days were fixed and stained to assess microvascular self-assembly. Microvascular network formation and extended branching morphology were observed in constructs regardless of print condition (no print, with gelatin, or with Pluronic F-127) (**Figure 6B, D, Figure S5B)**. Gelatin-printed constructs demonstrated a clear ring of void space where the gelatin had been evacuated from the construct upon incubation at 37 °C (**Figure 6Bi, 6Di)**. Some vascular formation and cell spreading was observed in areas coincident with the Pluronic F-127 structure print path (**Figure 6Bii, 6Dii)**. Quantification of vascular networks demonstrated no significant difference in vessel density between print and no print conditions (**Figure 6E**).

## 3. Discussion

Engineering functional and perfusable vasculature is critical towards engineering clinically relevant-sized tissue constructs. Hierarchical vasculature could provide the ideal support for constructs containing high densities of metabolically active cells through optimized transport of nutrients and waste. Strategies to create multiscale vasculature are beginning to show promise.^[9, 24, 42]^ In this work, we developed a novel approach that uses clickable PEGNB hydrogel microbeads as a suspension bath medium to support both bioprinting of mesoscale channels (within 50 – 1000 μm)^[3]^ with sacrificial bioinks and microvascular self-assembly within the interstitial voids between individual beads of secondarily-crosslinked granular constructs. These print-patterned, cell-laden constructs support the self-assembly of embedded endothelial cells into pericyte-invested, lumenized microvascular networks in the interstitial spaces between the RGD-functionalized PEGNB microbeads. Recent studies in the field have demonstrated the potential of individual elements of this strategy.^[39, 43]^ However, the combination of top-down suspension bioprinting, microbead photocrosslinking, and subsequent bottom-up microvascular self-assembly within a granular construct represents a novel and innovative platform. The use of PEGNB-based microbeads in this context is also novel as they may act as a blank slate for myriad customization. This platform makes substantial progress towards the creation of multiscale, hierarchical vasculature capable of supporting the metabolic demands of a wide-range of parenchymal cells in large tissue constructs approaching clinically relevant sizes (>1 cm^3^ in tissue volume).

Our approach exploits the fabrication of PEGNB hydrogel microbeads via microfluidic droplet generation, a relatively simple method of producing microbeads reliably and efficiently (PDI = 0.04) that does not require specialized equipment. PEGNB was chosen due to previous studies, including our own, demonstrating its ability to support microvascular self-assembly from cells embedded within the material,^[18, 44–46]^ as well as the capability to produce clickable PEGNB microbeads.^[47]^ The PEGNB beads were composed of a relatively high polymer weight percentage with low crosslinking ratio, leading to a high degree of swelling (∼1.83x). *In situ* dynamic rheology of the bead slurry in the presence of additional LAP and PEGDT demonstrated an increase in G’ when exposed to UV. This was indicative that the beads formed covalent inter-bead crosslinks, leading to a higher shear storage modulus for these granular constructs in comparison to the uncrosslinked state of the jammed bead slurry. The subsequent stability of G’ with prolonged UV exposure also indicates that the degree of crosslinking did not increase further, due to exhaustion of available LAP, PEGDT, and/or free norbornene arms on the beads.

Previous studies have demonstrated that clickable PEGNB beads are suitable as a jammed microgel ink,^[47]^ but their usage as a print bath medium suitable to support suspension bioprinting when jammed into a bead slurry as shown here is a novel application. Suspension bath printing requires a colloidal medium demonstrating Bingham plastic characteristics as a self-healing yield stress material.^[48]^ Mechanical characterizations were chosen based on existing suspension bath literature.^[10, 25, 27, 48–50]^ The PEGNB bead slurries exhibited shear thinning, strain yielding, and reversible solid-like and fluid-like behaviors. Decreasing viscosity with increasing shear rate indicated shear-thinning behavior of the colloidal PEGNB bead slurry, a necessary characteristic of suspension bath media to allow rapid and non-destructive movement of a printing needle or cone through the mixture. This was further supported by a strain sweep, where G’ was greater than G” for increasing strain magnitude, indicating solid-like behavior at lower strain. However, when the strain was increased beyond the critical strain (G” surpassed G’), the bead slurry demonstrated shear yielding behavior and a dominant fluid-like behavior. Jammed bead slurries oscillated between solid-like (G’ > G”) and fluid-like (G” > G’) behavior over the course of multiple cycles when subjected to low and high strain magnitudes, respectively. This confirmed that the transition between solid-like and fluid-like behavior was reversible and rapid (minimal thixotropy), as is necessary when performing suspension printing.

We demonstrated that the jammed bead slurry was supportive of suspension printing of soft bioink structures. Due to our goal to use bioprinting to template and pattern larger scale vascular structures in a granular microvascularized construct, we chose bioinks based on their sacrificial capability. Gelatin^[10, 25]^ and Pluronic F-127^[51–52]^ liquify at temperatures greater than 37 °C and less than 20 °C, respectively. The complex bioprinted structure was intentionally designed to traverse the full volume of the bead bath and provide a clear inlet/outlet to optimize evacuation of the sacrificial ink, thus producing a patterned void throughout the granular construct. We perfused the void with glycerol, demonstrating the suitability of the resulting channels (600 – 1000 μm diameter) to perfuse fluorescent dye, cells, and/or oxygenated media to characterize or support a vascularized construct in the future, similar to strategies previously demonstrated in the literature.^[6–7, 10, 12]^

Previous studies demonstrated the ability of granular materials to support spreading of mesenchymal stromal cells,^[31, 35, 47, 53–55]^ dermal fibroblasts,^[35–36]^ 3T3 fibroblasts,^[30]^ and neural stem cells.^[56]^ Strategies similar to ours to vascularize granular materials have recently been described. One demonstrated photoannealing of HUVEC-laden alginate-based microtissues and showed the formation of capillary-like networks between microtissues when collagen hydrogel was added to the inter-bead void space.^[40]^ A second combined an innovative surgical micropuncture technique to foster cellular infiltration of endothelial cells into GelMA-based granular hydrogels *in vivo*.^[39]^ Here, by embedding co-cultures of ECs and LFs in the inter-bead void space of the PEGNB granular constructs, we similarly observed robust microvascular self-assembly *in vitro*, but without the need for exogenous interstitial matrix. This allows us to harness the advantageous microporosity of granular scaffolds, such as aiding nutrient/waste diffusion, as well as the versatile PEG chemistry to functionalize the microbeads with additional biological cues. Furthermore, the ECs and LFs were embedded within the PEGNB microbead slurries prior to printing, then withstood printing and subsequent secondary crosslinking into granular constructs before robustly self-assembling into microvascular networks over the course of 7 days. While some spatial heterogeneity in the vascular networks was observed, this approach enabled vascularization of a large volume (>500 mm^3^).

We confirmed the presence of lumens within UEA-positive tubular structures and the expression of laminin β-1 and collagen IV, indicating network maturation through secretion of basement membrane proteins. While previous studies have focused on monoculture of ECs, the use of co-culture with stromal cells in this work supported vascular formation in our granular constructs. Perivascular association of LFs (or a subset of them) with the vascular networks was observed and then confirmed to express multiple pericyte markers, including the more restrictive NG2 and PDGFRβ, consistent with our previous studies.^[6]^ Although we did not vary the sizes of the microbead building blocks as part of this study, our swollen beads had diameters on the order of 400 µm, which was 2x-3x larger than those used in other studies involving granular hydrogels ^[35, 39]^. The use of relatively large microbeads, and their comparatively large pores, may have facilitated EC self-assembly into microvascular networks even in the absence of additional interstitial matrix in part by facilitating a 2.5D-like migration phenotype from the cells between microbeads.

Optimizing the LAP concentration was an important consideration towards achieving favorable cellular responses in photocrosslinked granular materials. Upon UV exposure, free radicals are generated^[57]^ to initiate the crosslinking between thiol and ene groups. Radicals are reactive and cytotoxic,^[58–59]^ and as such, the likely explanation for the poor cell morphology observed when higher concentrations of LAP were used. Cell density also supported cellular assembly up to a point of saturation, as cell morphology was improved when increased from 5×10^5^ cells/mL to 1×10^6^ cells/mL but there was no significant difference in vascular density between constructs containing 2×10^6^ cells/mL and 4×10^6^ cells/mL constructs. This may be due to faster nutrient depletion and waste production at higher cell densities limiting network formation when the volume of media exchanged and frequency of exchanges remains the same. Additionally, available inter-bead void space may have been limited due to the higher cell density. Notably, we achieved robust microvascular networks using nondegradable RGD-modified PEGNB beads, with cells interacting with the beads via the RGD and with other cells via the inter-bead voids. While not explored here, the use of degradable PEGNB beads or degradable inter-bead crosslinks in these constructs could facilitate remodeling both *in vitro* and *in vivo*, and may also influence the dependence of microvascular assembly on cell density since cells could also invade through the beads in addition to around them.

Finally, we showed that printing does not significantly affect microvascular self-assembly in the granular constructs. While cell viability, morphology, and behavior can be impacted by shear forces and mechanical disturbance,^[60–61]^ we observed no adverse effects from bioprinting into cell-laden PEGNB slurries. This may be attributed to the shear-thinning behavior of the jammed bead slurry, which may protect the cells from high shear stress as the print needle moves through the bath. Cell viabilities in granular materials were lower than in bulk PEG hydrogels of similar compositions; however, the bulk PEG gel controls had a total volume of 50 µL, while the granular constructs were 500 µL. This 10x difference in size may have contributed to the observed differences in cell viability, in addition to increased LAP concentration in the granular constructs. Lacking microporosity, it is possible that the bulk PEG gels would have impaired viability if made at a 500 µL volume due to diffusion limitations. More importantly, cell viability was not significantly different on day 1 of culture between print and no print conditions, and the number of viable cells in the granular constructs was still high enough to reliably support microvascular assembly, whether printed into or not. The viability and maintained ability of the cells to self-assemble into microvascular networks post-print and ink evacuation demonstrates this platform is supportive of complex multicellular morphogenesis, even with external manipulations to the system. As such, the platform is expected to support the growth of an endothelial monolayer within the print-patterned mesoscale channels via endothelial cell seeding, a critical next step to achieve multi-scale vasculature in our system.

## 4. Conclusion

Hierarchical vasculature is necessary to achieve tissue engineered constructs of clinically-relevant sizes. Towards the goal of creating such multiscale vasculature, we demonstrated the use of jammed PEGNB microbeads to support sacrificial patterning of mesoscale structures via suspension bioprinting in the presence of co-cultures of ECs and LFs, then subsequently clicked these microbeads together via thiol-ene photocrosslinking to support the self-assembly of microvasculature within the inter-bead voids over 7 days in culture. Clicked granular constructs maintained mechanical stability post-print and throughout cell culture. ECs formed interconnected vascular networks with hollow lumens, while LFs exhibited perivascular association and supported EC elongation and network formation. Collectively, our data show that suspension bioprinting and microvascular self-assembly were compatible and could occur within the same PEGNB granular bed. The porosity of the granular construct may also facilitate host tissue invasion if implanted *in vivo.* Future efforts will be focused on endothelialization of patterned void spaces within the microvascularized granular constructs to produce patent mesoscale vessel structures and characterization of the inosculation of these structures with the self-assembled microvasculature.

## 5. Experimental Section

### 5.1 PEG Microgel Fabrication

8-arm PEGNB (40 kDa; Creative PEGWorks, Durham, NC), 4-arm PEGNB (20 kDa; Creative PEGWorks, Durham, NC), and lithium phenyl-2,4,6-trimethylbenzoylphosphinate (LAP; Sigma-Aldrich, St. Louis, MO) were purchased from commercial sources that provide the percent substitution of norbornene by NMR and purity by HPLC, respectively. The thiol containing adhesive peptide Ac-CGRGDS-NH2 (RGD; AAPPTEC, Louisville, KY) which contains an N-terminal acetylation and a C-terminal amidation, was dissolved in 25 mM acetic acid, filtered through 0.22 µm filters (Sigma-Aldrich, St. Louis, MO), lyophilized for 48 hours and stored in a desiccator at −20 °C. The thiol content of the peptide aliquots was determined using Ellman’s reagent (Thermo Fisher, Waltham, MA). PEG-dithiol (PEG-DT, 3400 Da; Laysan Bio, Arab, AL) was used as a nondegradable crosslinker; PEG-DT purity was provided by the manufacturer.

To form sterile beads, 8-arm PEGNB, LAP, and PEGDT were suspended in phosphate-buffered saline (PBS, pH 7.4, 1X; Gibco, Waltham, MA) and sterile filtered through 0.22 µm filters. PEG hydrogel precursor solutions were made, consisting of 10 wt% PEGNB (w/v), 2 mM LAP, 2 mM RGD, PEGDT, and PBS. Crosslinking ratio was controlled via addition of PEGDT to achieve 37.5% crosslinking of available norbornene arms after accounting for RGD concentration. The precursor solution was then gently mixed by pipetting. Acellular, non-degradable poly(ethylene glycol)-norbornene (PEGNB) microbeads were formed from this precursor solution via microfluidic droplet generation using a polydimethylsiloxane (PDMS) flow-focusing device with channel dimensions of 150 x 150 μm before the junction and 150 x 200 μm after the junction. Sterile 0.5% perfluoropolyether (PFPE, 008-FluoroSurfactant; RAN Biotech, Beverly, MA) in NOVEC 7500 (3M, St. Louis, MO) was used to pinch the PEG precursor solution. Solutions were flowed into the device via syringe pumps (KD Scientific, Holliston, MA) at 15 µL/min (PEG precursor) and 30 µL/min (PFPE in NOVEC 7500). Beads were then collected in a sterile dish. After all PEG precursor was expelled, the beads were exposed to 6-Watt LED 365 nm Gooseneck Illuminator (AmScope, Feasterville, PA) and irradiated with UV for 5 min at 14.2 mW/cm^2^ to crosslink via thiol-ene photopolymerization. The beads were then isolated from the oil by removing the excess oil, rinsing with sterile 1% Pluronic F-68 (Thermo Fisher, Waltham, MA) twice and then PBS twice. The isolated beads were then swollen in PBS overnight.

### 5.2 Mechanical Characterization

Rheological properties of the PEGNB beads and PEGNB clicked granular constructs were measured using an AR-G2 rheometer (TA Instruments, New Castle, DE) with a 20 mm parallel plate measurement head. To measure the properties of the bead slurry for suspension bath printing applications, PEGNB microbeads were jammed via low pressure syringe-based vacuum filtration on 40 µm filters (pluriSelect, El Cajon, CA). Beads were then transferred onto the Peltier plate and centered under the measurement head. The head was then lowered to a 1 mm gap. To demonstrate shear-thinning properties, viscosity was measured with increasing shear rate (0.01 to 100 s^-1^). The beads were also subjected to strain ramp (0.01 to 100%, 1 Hz) to demonstrate strain-yielding properties. Shear recovery and reversible solid-like and fluid-like properties were demonstrated via oscillatory thixotropy tests, in which 1% and 150% strain were alternatingly applied at 1 Hz at 125 s intervals.

In-situ dynamic UV rheology using a UV LED accessory (TA Instruments, New Castle, DE) was performed to demonstrate the photopolymerization of the jammed bead slurry into a clicked granular construct. 500 µL of jammed beads were mixed with 10 µL of 12.5 mM LAP, 20 µL of 7.35 mM PEGDT, and 50 µL PBS to match the preparation of cell-laden granular constructs. This mixture was then placed on the 20 mm quartz plate with the UV emitting diodes. Shear storage modulus was measured over a time sweep performed with 1% strain amplitude and 1 rad/s frequency. At 40 s, the UV was turned on at an intensity of 13.2 mW/cm^2^ and stayed on for the remainder of the time sweep.

The shear storage moduli (G’) of the clicked granular constructs, acellular or cell-laden granular constructs, were measured on days 1 and 7 after incubation in EGM2. Acellular constructs were prepared in the same manner as the cell-laden constructs described below, but without cells. EGM2 was changed in acellular constructs in the same manner as cell-laden constructs. On day 1 or day 7, constructs were removed from the well plate and centered between the Peltier plate and measurement head (covered with P800 sandpaper). Shear storage modulus was averaged over a 1 min time sweep measured at 37 °C, 5% strain amplitude, 1 rad/s frequency, and a normal force of 0.05 N.

### 5.3 Imaging-based Characterization

Bead diameter and void fraction were characterized by incubating non-fluorescent beads or clicked granular constructs made of non-fluorescent beads with 5 mg/mL TRITC-dextran (155 kDa; Sigma-Aldrich, St. Louis, MO)^[62]^ for 15 minutes. Constructs and beads were then imaged using an Olympus IX81 microscope equipped with a disc scanning unit (DSU; Olympus America, Center Valley, PA) and Metamorph Premier software (Molecular Devices, Sunnyvale, CA). Bead diameter and percent porosity were measured and calculated via analysis using Fiji^[63]^. Polydispersity index (PDI) was calculated as (standard deviation/mean)^2^ of bead diameters across multiple batches.

To visualize clicked granular constructs and void space, 10 wt% 8-arm PEGNB beads with 37.5% crosslinking ratio and 2 mM thiolated rhodamine B (1K; BiochemPEG, Watertown, MA) were formed. Beads were formed into photopolymerized granular constructs by mixing jammed beads with 12.5 mM LAP and 7.35 mM PEGDT and UV crosslinking for 1 minute. Clicked granular constructs were then incubated with 5 mg/mL FITC-dextran (2000 kDa; Sigma-Aldrich, St. Louis, MO) and imaged using a Zeiss LSM800 (Zeiss, Oberkochen, Germany) laser scanning confocal microscope at slice intervals of 3.88 µm over a total stack thickness of 700 µm. Porosity data was analyzed using the Analyze Particles module in Fiji and averaged over 8 regions of interest across multiple granular constructs.

### 5.4 Cell Culture

Human umbilical vein endothelial cells (ECs) were isolated from fresh umbilical cords (obtained via an IRB-exempt protocol from the University of Michigan Mott’s Children’s Hospital), cultured in fully supplemented EGM2 (Lonza, Inc., Walkersville, MD), and used until passage 4. Normal human lung fibroblasts (LFs; Lonza) were cultured in high glucose Dulbecco’s modified eagle medium (DMEM; Gibco, Waltham, MA) supplemented with 10% fetal bovine serum (FBS; Gibco) and used until passage 9. EGM2 and DMEM were additionally supplemented with 1X Antibiotic-Antimycotic (Gibco). All cells were cultured at 37 °C and 5% CO_2_ with medium exchanges every other day.

### 5.5 Cell-laden Granular and Bulk PEG Gel Preparation and Characterization

10 wt% 8-arm PEGNB 37.5% crosslinked beads with 2 mM RGD were used to make cell-laden granular constructs. Beads were jammed via low pressure vacuum filtration as described above, and then 500 µL of beads were transferred to a well of a non-tissue culture treated 24-well plate. A non-tissue culture treated plate was used to minimize culture of cells on the plastic rather than in the granular construct. A cell pellet of ECs and LFs in a 1:1 EC:LF ratio was resuspended in serum-free EGM2 media. Small volumes of sterile-filtered LAP (10 µL, 12.5 mM) and PEGDT (20 µL, 7.35 mM) suspended in serum-free EGM2 were then added to the 500 µL jammed bead volume. 50 µL of the EC and LF (EC-LF) cell suspension in serum-free EGM2 was then added to the bead volume to achieve a final cell density of 2×10^6^ cells/mL or 4×10^6^ cells/mL and mixed briefly with a sterile pipette tip. The constructs were then placed 5 inches under a 6-Watt LED 365 nm Gooseneck Illuminator and irradiated with UV at max intensity for 1 minute, corresponding to 13.2 mW/cm^2^, to crosslink the beads into a granular construct. 2 mL of warmed EGM2 was then immediately added to the well and changed on day 1 of culture and every 2 days afterwards.

To form cell-laden bulk gels, a dithiol-containing crosslinking peptide containing two matrix metalloproteinase-(MMP-) sensitive cleavage sites, Ac-GCRDVPMS↓MRGGGVPMS↓MRGGDRCG-NH2 (“dVPMS”, cleavage sites indicated by ↓; AAPPTEC, Louisville, KY), was first dissolved in 25 mM acetic acid, filtered through 0.22 µm filters, lyophilized for 48 hours, and stored in a desiccator at −20 °C. The thiol content of the peptide aliquots was determined using Ellman’s reagent. Bulk PEG hydrogel precursor solutions composed of 3 wt% 4-arm PEGNB with 60% dVPMS crosslinking (0.6 thiols per norbornene after accounting for RGD concentration), 1 mM RGD, and 1 mM LAP were produced as previously described.^[18]^ Hydrogel precursor solutions containing cells (2×10^6^ cells/mL EC-LF in a 1:1 ratio) were gently vortexed before 50 µL was pipetted into a 1 mL syringe with the needle end cut off to cast the gel. Hydrogels were then irradiated under UV for 90 s at 50 mW/cm^2^. Gels were then expelled out of the syringes into 24-well plates containing 2 mL of EGM2, with the media changed on day one and every two days afterwards.

To characterize viability of cells in the cell-laden granular or bulk constructs, gels and constructs were stained with a LIVE/DEAD cell imaging kit (Invitrogen, Carlsbad, CA) on day one of culture. Calcein AM and BOBO-3 iodide were used to visualize live and dead/dying cells, respectively. 300 µm confocal z-stacks (50 µm/slice, 7 slices/stack) were acquired and collapsed into maximum intensity projections prior to analysis using Fiji. Percent viability was calculated and normalized to the average of the no print condition for statistical analysis.

To characterize microvascular assembly, cell-laden granular constructs were fixed on day 7 with zinc formalin (Z-fix; Anatech, Battle Creek, MI) for 15 min, then washed three times with PBS for 5 min. Constructs were then stained overnight with rhodamine-conjugated lectin from *Ulex europaeus* agglutinin I (UEA, 1:200; Vector Labs, Newark, CA), 4’, 6-diamidino-2 phenylindol (DAPI, 1 ug/mL; Thermo Fisher, Waltham, MA), and AlexaFluor 488 phalloidin (1:100; Thermo Fisher, Waltham, MA) to label ECs, cell nuclei, and F-actin, respectively. Stain was removed from the samples before washing once with PBS for 5 min and overnight with PBS prior to imaging. Samples were then removed from well-plates and placed on glass slides to image. 300 µm confocal z-stacks (50 µm/slice, 7 slices/stack) were then acquired using the Olympus IX81 DSU and collapsed into maximum intensity projections using Fiji prior to analysis. Bulk PEG gels were processed similarly, but cut in half using a razor blade prior to staining^[18]^ and imaged on the flat cut side. Vascular densities were quantified on 4X magnification stacks using the Angiogenesis Tube Formation module in Metamorph and reported as vessel density length per volume. At least three regions of interest were quantified per construct and averaged. Images of full constructs were taken via the scan slide module in Metamorph at 4X magnification with the Olympus IX81 DSU.

### 5.6 Immunofluorescent Staining and Imaging

For immunofluorescent staining of basement membrane proteins and pericyte markers, constructs were fixed with Z-fix at day 7. A stock solution of 10X Tris buffered saline (TBS) was prepared with 44 g NaCl (Thermo Fisher, Waltham, MA), 15.75 g Tris (Bio-Rad, Hercules, CA), and 500 mL ddH2O and adjusted to pH 7.4. Granular constructs were permeabilized in 0.5% Triton X-100 (Thermo Fisher, Waltham, MA) in 1X TBS adjusted to pH 7.6 for 1 hour at room temperature and rinsed 3 times for 1 hour in 0.1% Tween-20 (Thermo Fisher, Waltham, MA) in 1X TBS adjusted to pH 7.6 (TBS-T). Constructs were blocked in antibody diluting (AbDil) solution (2% bovine serum albumin in TBS-T) at 4 °C overnight. Constructs were then incubated overnight at 4 °C with primary antibodies for collagen IV (1:200, rabbit IgG; Abcam, Waltham, MA) or laminin β-1 (1:200, rabbit IgG; Invitrogen, Carlsbad, California) or alpha-smooth muscle actin (1:200, mouse monoclonal; Sigma Aldrich, St. Louis, MO) or neuron-glial antigen 2 (NG2, 1:200, rabbit monoclonal recombinant; Abcam, Waltham, MA) or platelet derived growth factor receptor β (PDGFR-β, 1:100, rabbit polyclonal; Sigma Aldrich, St. Louis, MO) diluted in AbDil solution. Constructs were then rinsed 3 times for 1 hour with TBS-T before being stained with UEA, DAPI, and either AlexaFluor 488 goat anti-rabbit (1:200, IgGH+L; Thermo Fisher, Waltham, MA) or AlexaFluor 488 goat anti-mouse (1:200, IgGH+L; Thermo Fisher, Waltham, MA) diluted in AbDil solution. The constructs were incubated with the secondary staining solution overnight at 4 °C, and then rinsed 3 times for 1 hr in TBS-T prior to imaging. Representative images were taken at 4X and 10X magnification using the Olympus IX81 DSU and shown as maximum intensity projections of 300 µm stacks (50 µm/slice, 7 slices/stack).

### 5.7 Printing

Bioprinting was performed using an Allevi 3 (Allevi by 3D Systems, Philadelphia, PA). All inks were extruded using 30G, ½ inch blunt metal needle tips (BSTEAN). Sterilized needle tips were used when printing into cell-laden bead baths. All bioprinting took place inside a biosafety cabinet to maintain sterility. Computer aided-design (CAD) structures were designed with SolidWorks (Dassault Systemes, France) and TinkerCad (Autodesk, San Francisco, CA). 40% (w/v) Pluronic F-127 (Sigma Aldrich, St. Louis, MO) and 2.5% (w/v) gelatin (Thermo Fisher, Waltham, MA) were prepared in PBS to be used as sacrificial bioinks. Food dye (Kroger, Cincinnati, Ohio) was used to visualize the inks. For printing into cell-laden bead baths, bioinks were sterile filtered with 0.22 µm syringe filters. Inks were loaded into 5 mL syringes (Allevi by 3D Systems, Philadelphia, PA) to prepare for printing.

For gelatin printing, the ink-filled syringe was placed in the printer and the cartridge was set to 50°C for 10 min to melt the gelatin. After melting, the cartridge was cooled at 15 °C for 25 min to reach the gelation level necessary for printing. The optimized extrusion pressure was 50 PSI with a print speed of 6 mm/s. For Pluronic F-127 printing, the ink-filled syringe was placed in the printer without temperature control, with an optimized extrusion pressure of 80 PSI and print speed of 6 mm/s.

To prepare the beads for suspension bath printing, jammed bead slurries were prepared as described above. A PDMS mold (20 mm diameter x 15 mm height) was then placed on a glass slide to act as a well for printing. 1.5 mL to 2 mL of beads were then added to the well. To prepare beads for crosslinking post-print, 10 µL of 100 mM LAP in PBS and 20 µL of 58.8 mM PEGDT in PBS were added to the jammed slurry prior to printing and mixed well.

To produce acellular granular constructs with sacrificed voids for perfusion, gelatin structures were printed into the jammed bead slurry with LAP and PEGDT. After printing, the beads were UV photopolymerized for 5 min at 37.5 mW/cm^2^. Afterwards, the PDMS mold could be removed from the glass slide and the construct with embedded print was free standing. The granular construct was then moved to a container of warmed PBS (>37 °C) to melt and evacuate the gelatin for 20 min. Afterwards, the construct was ready for perfusion or storage in PBS long-term. Glycerol (Sigma Aldrich, St. Louis, MO) was used to perfuse the print-patterned voids within the granular constructs. Visualized with the addition of food dye, a syringe pump was used to flow glycerol into the construct at a rate of 30 µL/min using a 20G, ½ in blunt tip needle.

To print into cell-laden bead baths, the cell-laden bead slurry was prepared as described above (Experimental Section 5.5). Prior to the final step of UV crosslinking, the well plate was moved to the printer, where sterile gelatin or Pluronic F-127 were printed in the shape of a cylinder (8 mm diameter x 1 mm height) into the cell-laden bead bath. After printing, the constructs were UV photopolymerized for 1 min at 13.2 mW/cm^2^, before warmed EGM2 was added. Media was changed the next day and every 2 days afterwards until day 7.

### 5.8 Statistics

Statistical analysis was performed using Prism (GraphPAD, La Jolla, CA). Data are presented as mean +/-standard deviation of at least three independent experimental replicates. For independent replicates in each condition, data from two technical replicates was averaged together, resulting in one data point per independent replicate. Data was analyzed via unpaired t-test when comparing only two groups or one-way ANOVA with Tukey post-hoc testing when comparing more than two groups. A value of alpha ≤ 0.05 was considered significant.

## 6. Acknowledgments

Research reported in this publication was supported by the National Heart, Lung, and Blood Institute of the National Institutes of Health under Award Number R01-HL085339 and the Leland Professorship endowment. IWZ was partially supported by the Cellular Biotechnology Training Program at the University of Michigan (T32-GM145304). NEF was partially supported by the Tissue Engineering and Regeneration Training Program at the University of Michigan (T32-DE007057) and the Rackham Merit Fellowship. AJM was partially supported by the Training Program in Translational Cardiovascular Research and Entrepreneurship at the University of Michigan (T32-HL125242). BMB and FSM acknowledge support from NIH (R01-EB030474), the National Science Foundation (CELL-MET ERC, EEC-1647837), and a Scientific Research Initiative grant from the Biosciences Initiative of the University of Michigan. The content is solely the responsibility of the authors and does not necessarily represent the official views of the National Institutes of Health or the National Science Foundation. Some assets in figures were created in BioRender (licensed to A. Putnam).

## References

[1] M. A. Traore, S. C. George, Tissue Eng Part B Rev 2017, 23, 505.

[2] R. K. Jain, P. Au, J. Tam, D. G. Duda, D. Fukumura, Nat Biotechnol 2005, 23, 821.

[3] E. A. Margolis, N. E. Friend, M. W. Rolle, E. Alsberg, A. J. Putnam, Trends Biotechnol 2023, 41, 1400.

[4] X. Meng, Y. Xing, J. Li, C. Deng, Y. Li, X. Ren, D. Zhang, Front Cell Dev Biol 2021, 9, 639299.

[5] C. O’Connor, E. Brady, Y. Zheng, E. Moore, K. R. Stevens, Nat Rev Mater 2022, 7, 702.

[6] E. A. Margolis, D. S. Cleveland, Y. P. Kong, J. A. Beamish, W. Y. Wang, B. M. Baker, A. J. Putnam, Lab Chip 2021, 21, 1150.

[7] L. Debbi, B. Zohar, M. Shuhmaher, Y. Shandalov, I. Goldfracht, S. Levenberg, Biomaterials 2022, 280, 121286.

[8] H. G. Song, A. Lammers, S. Sundaram, L. Rubio, A. X. Chen, L. Li, J. Eyckmans, S. N. Bhatia, C. S. Chen, Adv Funct Mater 2020, 30.

[9] B. Zhang, M. Montgomery, M. D. Chamberlain, S. Ogawa, A. Korolj, A. Pahnke, L. A. Wells, S. Masse, J. Kim, L. Reis, A. Momen, S. S. Nunes, A. R. Wheeler, K. Nanthakumar, G. Keller, M. V. Sefton, M. Radisic, Nat Mater 2016, 15, 669.

[10] M. A. Skylar-Scott, S. G. M. Uzel, L. L. Nam, J. H. Ahrens, R. L. Truby, S. Damaraju, J. A. Lewis, Science Advances 2019, 5, eaaw2459.

[11] B. Grigoryan, S. J. Paulsen, D. C. Corbett, D. W. Sazer, C. L. Fortin, A. J. Zaita, P. T. Greenfield, N. J. Calafat, J. P. Gounley, A. H. Ta, F. Johansson, A. Randles, J. E. Rosenkrantz, J. D. Louis-Rosenberg, P. A. Galie, K. R. Stevens, J. S. Miller, Science 2019, 364, 458.

[12] I. S. Kinstlinger, G. A. Calderon, M. K. Royse, A. K. Means, B. Grigoryan, J. S. Miller, Nat Protoc 2021, 16, 3089.

[13] J. Son, S. J. Hong, J. W. Lim, W. Jeong, J. H. Jeong, H. W. Kang, Small Methods 2021, 5, e2100632.

[14] J. S. Miller, K. R. Stevens, M. T. Yang, B. M. Baker, D. H. Nguyen, D. M. Cohen, E. Toro, A. A. Chen, P. A. Galie, X. Yu, R. Chaturvedi, S. N. Bhatia, C. S. Chen, Nat Mater 2012, 11, 768.

[15] A. Lee, A. R. Hudson, D. J. Shiwarski, J. W. Tashman, T. J. Hinton, S. Yerneni, J. M. Bliley, P. G. Campbell, A. W. Feinberg, Science 2019, 365, 482.

[16] X. Chen, A. S. Aledia, C. M. Ghajar, C. K. Griffith, A. J. Putnam, C. C. Hughes, S. C. George, Tissue Eng Part A 2009, 15, 1363.

[17] N. E. Friend, A. Y. Rioja, Y. P. Kong, J. A. Beamish, X. Hong, J. C. Habif, J. R. Bezenah, C. X. Deng, J. P. Stegemann, A. J. Putnam, Sci Rep 2020, 10, 15562.

[18] N. E. Friend, A. J. McCoy, J. P. Stegemann, A. J. Putnam, Biomaterials 2023, 295, 122050.

[19] S. Kim, H. Lee, M. Chung, N. L. Jeon, Lab Chip 2013, 13, 1489.

[20] A. N. Stratman, K. M. Malotte, R. D. Mahan, M. J. Davis, G. E. Davis, Blood 2009, 114, 5091.

[21] E. M. Meijer, C. G. M. van Dijk, R. Kramann, M. C. Verhaar, C. Cheng, Tissue Eng Part B Rev 2022, 28, 1.

[22] E. Warren, S. Gerecht, Vasc Biol 2023, 5.

[23] F. Helms, S. Zippusch, J. Theilen, A. Haverich, M. Wilhelmi, U. Boer, Biotechnol Bioeng 2022, 119, 2239.

[24] A. A. Szklanny, M. Machour, I. Redenski, V. Chochola, I. Goldfracht, B. Kaplan, M. Epshtein, H. Simaan Yameen, U. Merdler, A. Feinberg, D. Seliktar, N. Korin, J. Jaros, S. Levenberg, Adv Mater 2021, 33, e2102661.

[25] T. J. Hinton, Q. Jallerat, R. N. Palchesko, J. H. Park, M. S. Grodzicki, H.-J. Shue, M. H. Ramadan, A. R. Hudson, A. W. Feinberg, Science Advances 2015, 1, e1500758.

[26] O. Jeon, Y. B. Lee, H. Jeong, S. J. Lee, D. Wells, E. Alsberg, Mater Horiz 2019, 6, 1625.

[27] A. M. Compaan, K. Song, W. Chai, Y. Huang, ACS Appl Mater Interfaces 2020, 12, 7855.

[28] A. McCormack, C. B. Highley, N. R. Leslie, F. P. W. Melchels, Trends Biotechnol 2020, 38, 584.

[29] A. C. Daly, L. Riley, T. Segura, J. A. Burdick, Nat Rev Mater 2020, 5, 20.

[30] C. B. Highley, K. H. Song, A. C. Daly, J. A. Burdick, Adv Sci (Weinh) 2019, 6, 1801076.

[31] S. Xin, O. M. Wyman, D. L. Alge, Adv Healthc Mater 2018, 7, e1800160.

[32] J. E. Mealy, J. J. Chung, H. H. Jeong, D. Issadore, D. Lee, P. Atluri, J. A. Burdick, Adv Mater 2018, 30, e1705912.

[33] A. E. Widener, M. Bhatta, T. E. Angelini, E. A. Phelps, Biomater Sci 2021, 9, 2480.

[34] D. R. Griffin, M. M. Archang, C. H. Kuan, W. M. Weaver, J. S. Weinstein, A. C. Feng, A. Ruccia, E. Sideris, V. Ragkousis, J. Koh, M. V. Plikus, D. Di Carlo, T. Segura, P. O. Scumpia, Nat Mater 2021, 20, 560.

[35] D. R. Griffin, W. M. Weaver, P. O. Scumpia, D. Di Carlo, T. Segura, Nat Mater 2015, 14, 737.

[36] N. F. Truong, E. Kurt, N. Tahmizyan, S. C. Lesher-Perez, M. Chen, N. J. Darling, W. Xi, T. Segura, Acta Biomater 2019, 94, 160.

[37] L. Riley, L. Schirmer, T. Segura, Curr Opin Biotechnol 2019, 60, 1.

[38] L. J. Pruett, A. L. Taing, N. S. Singh, S. M. Peirce, D. R. Griffin, Acta Biomater 2022, 148, 171.

[39] Z. Ataie, S. Horchler, A. Jaberi, S. V. Koduru, J. C. El-Mallah, M. Sun, S. Kheirabadi, A. Kedzierski, A. Risbud, A. Silva, D. J. Ravnic, A. Sheikhi, Small 2024, 20, e2307928.

[40] M. Schot, M. Becker, C. A. Paggi, F. Gomes, T. Koch, T. Gensheimer, C. Johnbosco, L. P. Nogueira, A. van der Meer, A. Carlson, H. Haugen, J. Leijten, Adv Mater 2023, e2308949.

[41] J. M. Hughes, P. M. Budd, A. Grieve, P. Dutta, K. Tiede, J. Lewis, Journal of Applied Polymer Science 2015, 132.

[42] P. Wu, H. Asada, M. Hakamada, M. Mabuchi, Adv Mater 2023, 35, e2209149.

[43] Y. Fang, Y. Guo, B. Wu, Z. Liu, M. Ye, Y. Xu, M. Ji, L. Chen, B. Lu, K. Nie, Z. Wang, J. Luo, T. Zhang, W. Sun, Z. Xiong, Adv Mater 2023, 35, e2205082.

[44] M. R. Zanotelli, H. Ardalani, J. Zhang, Z. Hou, E. H. Nguyen, S. Swanson, B. K. Nguyen, J. Bolin, A. Elwell, L. L. Bischel, A. W. Xie, R. Stewart, D. J. Beebe, J. A. Thomson, M. P. Schwartz, W. L. Murphy, Acta Biomater 2016, 35, 32.

[45] A. Brown, H. He, E. Trumper, J. Valdez, P. Hammond, L. G. Griffith, Biomaterials 2020, 243, 119921.

[46] M. Sofman, A. Brown, L. G. Griffith, P. T. Hammond, Biomaterials 2021, 264, 120231.

[47] S. Xin, D. Chimene, J. E. Garza, A. K. Gaharwar, D. L. Alge, Biomater Sci 2019, 7, 1179.

[48] M. E. Cooke, D. H. Rosenzweig, APL Bioeng 2021, 5, 011502.

[49] A. M. Compaan, K. Song, Y. Huang, ACS Appl Mater Interfaces 2019, 11, 5714.

[50] M. E. Prendergast, J. A. Burdick, Adv Healthc Mater 2022, 11, e2101679.

[51] Y. Xu, Y. Hu, C. Liu, H. Yao, B. Liu, S. Mi, Materials (Basel) 2018, 11.

[52] B. Ren, K. Song, A. R. Sanikommu, Y. Chai, M. A. Longmire, W. Chai, W. L. Murfee, Y. Huang, Appl Phys Rev 2022, 9, 011408.

[53] A. S. Caldwell, G. T. Campbell, K. M. T. Shekiro, K. S. Anseth, Adv Healthc Mater 2017, 6.

[54] C. E. Miksch, N. P. Skillin, B. E. Kirkpatrick, G. K. Hach, V. V. Rao, T. J. White, K. S. Anseth, Small 2022, 18, e2200951.

[55] S. Xin, C. A. Gregory, D. L. Alge, Acta Biomater 2020, 101, 227.

[56] C. D. Morley, E. A. Ding, E. M. Carvalho, S. Kumar, Adv Mater 2023, 35, e2304212.

[57] B. D. Fairbanks, M. P. Schwartz, C. N. Bowman, K. S. Anseth, Biomaterials 2009, 30, 6702.

[58] S. J. Bryant, C. R. Nuttelman, K. S. Anseth, J Biomater Sci Polym Ed 2000, 11, 439.

[59] J. P. Kehrer, Crit Rev Toxicol 1993, 23, 21.

[60] S. Han, C. M. Kim, S. Jin, T. Y. Kim, Biofabrication 2021, 13.

[61] J. Hua, L. E. Erickson, T. Y. Yiin, L. A. Glasgow, Crit Rev Biotechnol 1993, 13, 305.

[62] T. H. Qazi, V. G. Muir, J. A. Burdick, ACS Biomater Sci Eng 2022, 8, 1427.

[63] J. Schindelin, I. Arganda-Carreras, E. Frise, V. Kaynig, M. Longair, T. Pietzsch, S. Preibisch, C. Rueden, S. Saalfeld, B. Schmid, J. Y. Tinevez, D. J. White, V. Hartenstein, K. Eliceiri, P. Tomancak, A. Cardona, Nat Methods 2012, 9, 676.

