## Supplemental Figures Document for "Clickable PEG-norbornene microgels support suspension bioprinting and microvascular assembly": Zhang, et al. PEG Bead Paper Supporting Information.docx

Movie S1: 3D animated video of void space and beads

Movie S2: Printing into bead bath

Movie S3: Perfusion into print patterned void – 3D squiggle structure

Movie S4: Perfusion into print patterned void – horizontal capillary bed structure

Movie S5: Lumen animation 1

Movie S6: Lumen animation 2

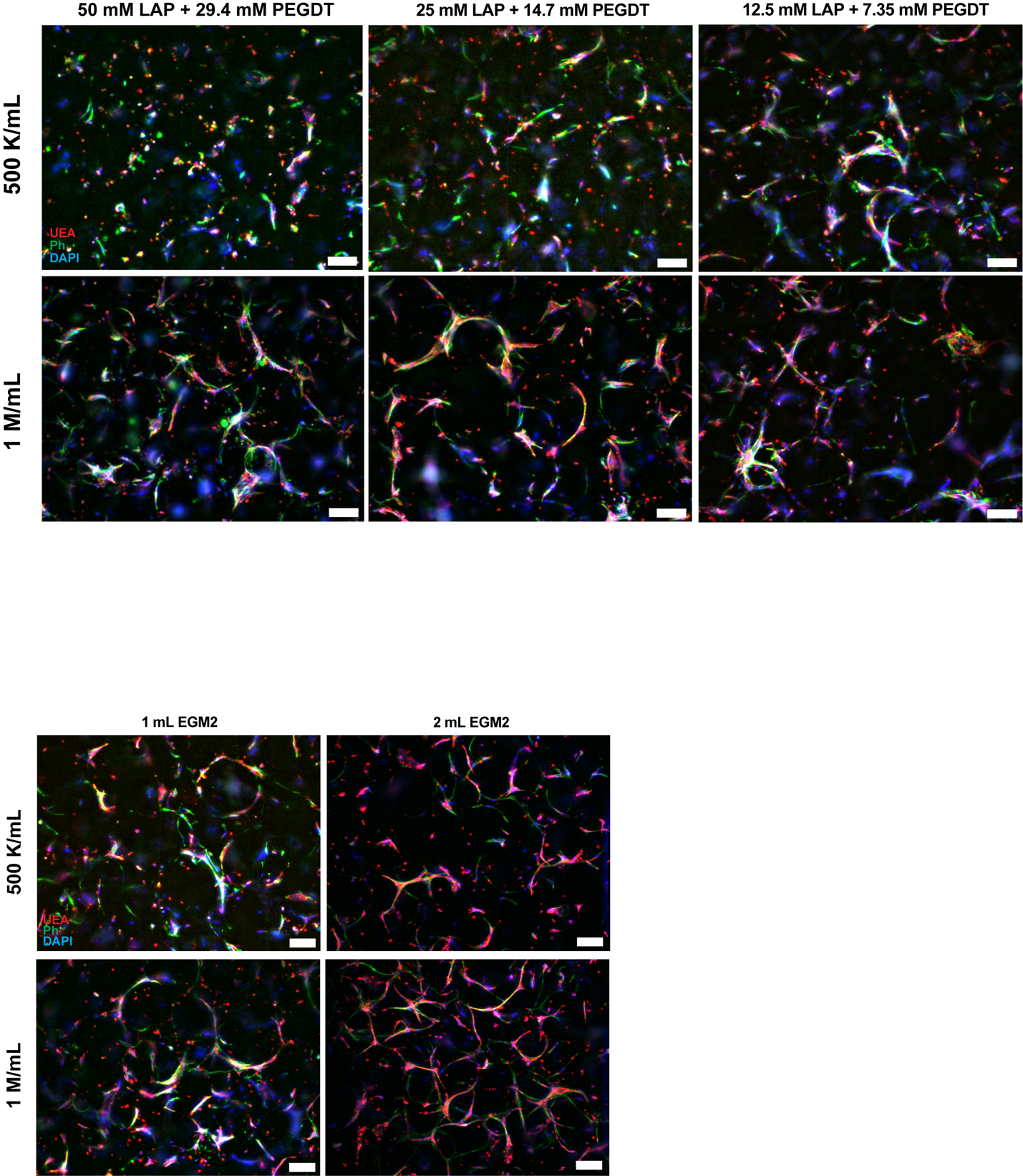

**Figure S1. Concentration of photoinitiator and crosslinker for secondary crosslinking of clickable microbeads impacts cell morphology.** Representative fluorescent microscopy images of cell morphology after 7 days of culture with 10 wt% PEG8NB 37.5% crosslinked 2 mM RGD granular constructs. Concentration of LAP and PEGDT were varied in addition to initial cell seeding density of a 1:1 ratio of HUVECs and NHLFs. Constructs were fed 1 mL of EGM2 per media change. Images taken at 4X magnification (scale bar = 200 µm; red = UEA, green = phalloidin, blue = DAPI).

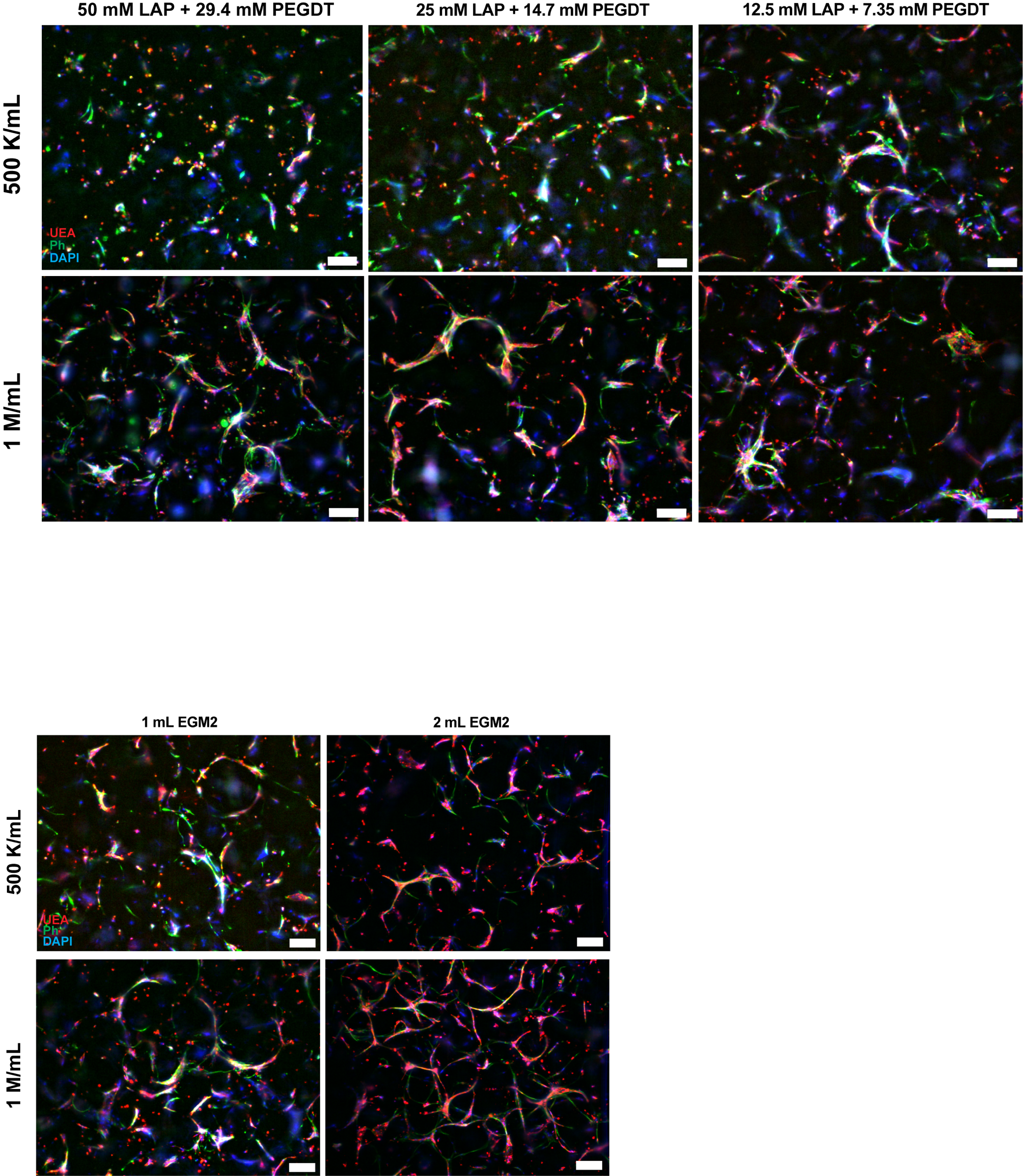

**Figure S2. Media volume impacts degree of microvascular self-assembly.** Representative fluorescent microscopy images of cell morphology in 10% PEG8NB 37.5% crosslinked 2 mM RGD granular constructs fed with 1 or 2 mL of EGM2 over the course of 7 days of culture. Constructs were crosslinked with the addition of 12.5 mM LAP and 7.35 mM PEGDT. Images taken at 4X magnification (scale bar = 200 µm; red = UEA, green = phalloidin, blue = DAPI).

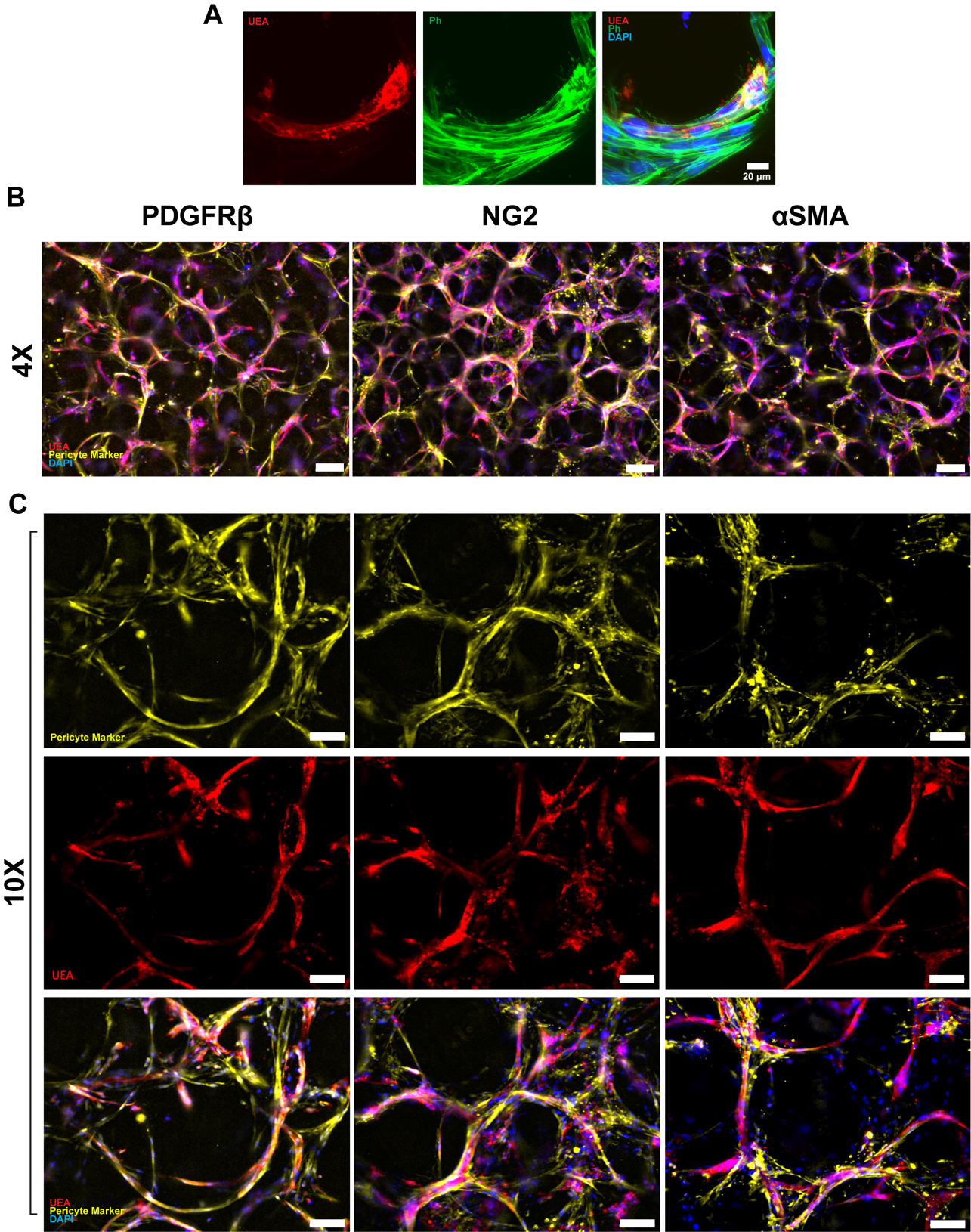

**Figure S3. Pericyte markers PDGFRβ, NG2, and αSMA exhibit close, sheath-like association with vascular projections.** Representative fluorescent images of EC-LF-laden granular constructs (2x10^6^ cells/mL) after 7 days of culture stained with A) phalloidin (green), UEA (red), and DAPI (blue) at 40X magnification, demonstrating fibroblast (UEA-negative, phalloidin-positive cell) association with tubular endothelial cell. Constructs were stained for PDGFRβ, NG2, or αSMA and imaged at B) 4X (200 µm scale bar) and C) 10X magnification (100 µm scale bar).

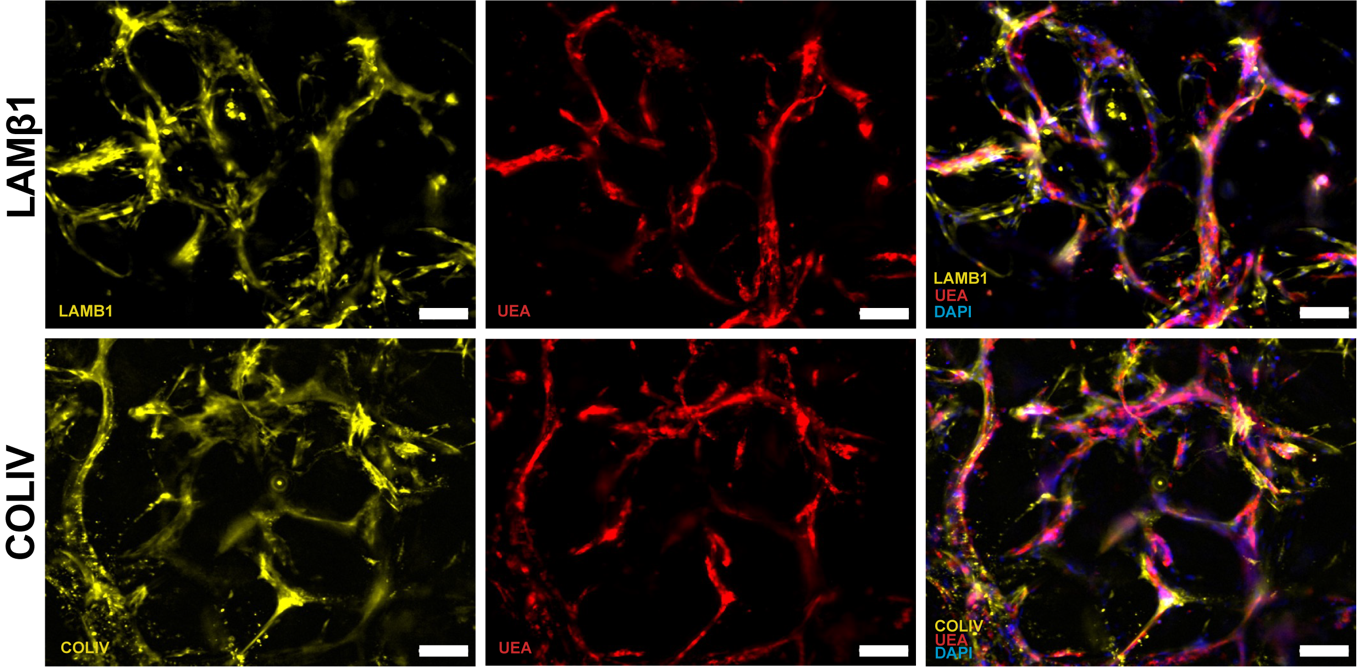

**Figure S4. Vascular projections are co-localized with basement membrane proteins, indicating mature vascular development.** Representative fluorescent images of EC-LF-laden granular constructs (2x10^6^ cells/mL) after 7 days of culture stained with laminin β-1 and collagen IV at 10X magnification (scale bar = 100 µm).

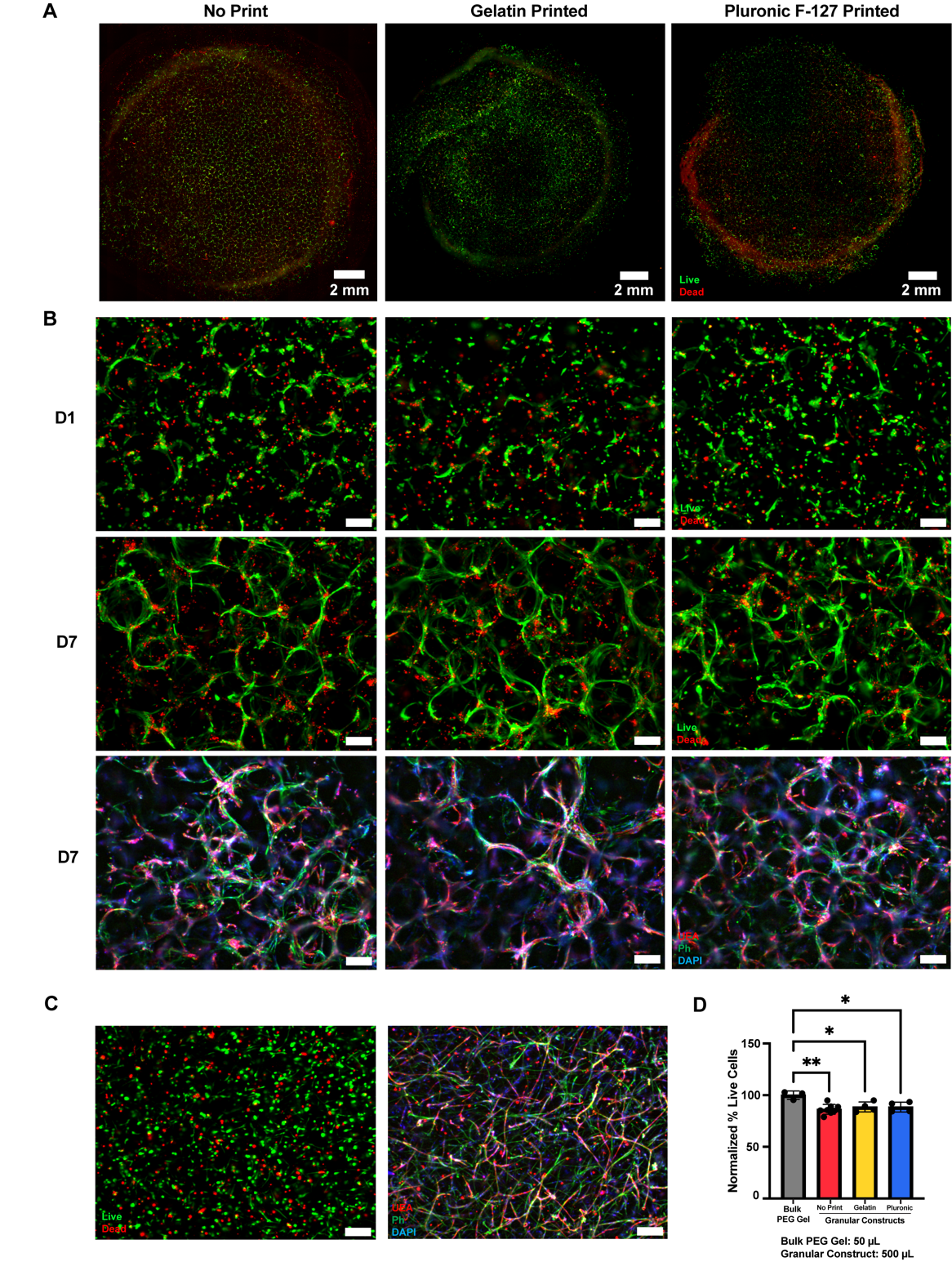

**Figure S5. Cell-laden constructs with or without printing on day 0 exhibited no significant difference in viability.** A) Representative fluorescent 4X scan-slide images of EC-LF-laden granular constructs (2x10^6^ cells/mL) on day 1, stained with live/dead indicator. B) Representative maximum intensity projection images (300 µm stack, 7 slice/stack) of EC-LF-laden granular constructs (2x10^6^ cells/mL) stained with live/dead indicator on day 1 and day 7 (red = dead, green = live) and microvascular self-assembly stained after 7 days of culture (red = HUVECs, green = f-actin, blue = nuclei) (4X magnification = 200 µm scale bar). C) EC-LF-laden bulk PEG gels (2x10^6^ cells/mL) were live/dead stained on day 1 and stained on day 7 for microvascular formation and imaged at 4X (scale bar = 200 µm). D) Viability of cells in cell-laden constructs on day 1 was analyzed via live/dead stain. Percent viability of no print granular constructs (n = 8), gelatin printed granular constructs (n = 4), and Pluronic F-127 printed granular constructs (n = 4) when normalized to the bulk PEG gel average (n = 3). *: p ≤ 0.05, **: p ≤ 0.01
